# Infection of *Plasmodiophora brassicae* changes the fungal endophyte community of tumourous stem mustard roots as revealed by high-throughput sequencing and culture-dependent methods

**DOI:** 10.1101/590018

**Authors:** Xueliang Tian, Diandong Wang, Zhenchuan Mao, Limei Pan, Jingjing Liao, Zhaoming Cai

**Author notes:** Corresponding author: Diandong Wang.

## Abstract

Diverse fungal endophytes live in plants and are shaped by some abiotic and biotic stresses. Plant disease as particular biotic stress possibly gives an impact on the communities of fungal endophytes. In this study, clubroot disease caused by an obligate biotroph protist, *Plasmodiophora brassicae*, was considered to analyze its influence on the fungal endophyte community using an internal transcribed spacer (ITS) through high-throughput sequencing and culture-dependent methods. The results show that the diversity of the endophyte community in the healthy roots was much higher than the clubroots. Ascomycota was the dominant group of endophytes (including *Phoma, Mortierella, Penicillium*, etc.) in the healthy roots while *P*. *brassicae* was the dominant taxon in the clubroots. Hierarchical clustering, principal component analysis (PCA) and principal coordinates analysis (PCoA) indicated significant differences between the endophyte communities in the healthy roots and clubroots. Linear discriminant analysis effect size (LefSe) analysis showed that the dominant genera could be regarded as potential biomarkers. The endophyte community in the healthy roots had a more complex network compared with the clubroots. Also, many plant pathogenic *Fusarium* were isolated from the clubroots by the culture-dependent method. The outcome of this study illustrates that *P*. *brassicae* infection may change the fungal endophyte community associated with the roots of tumourous stem mustard and facilitates the entry of soil pathogen into the roots.

## Introduction

Fungal endophytes have a close relationship with the host plants and they live in their tissues [1,2]. They provide many ecological and physiological advantages to their hosts, such as growth promotion [3], resistance to plant pathogens and adaptability to various abiotic stresses such as temperature, pH, and osmotic pressure as well as biotic stresses [4-6]. Similarly, the fungal endophyte communities are also affected by abiotic and biotic stresses [7]. Plant disease as particular biotic stress results in significant changes in the physiology of the plant. These changes and pathogen itself may affect the diversity and composition of the fungal endophyte inhabiting the plant tissues and their interactions with their host plant [8-10].

Clubroot is a severe disease of cruciferous crops such as cabbage and cauliflower, which is caused by the biotrophic *Plasmodiophora brassicae* Woronin [11]. The infected root cells undergo abnormal cell division and enlargement resulting in the formation of spindle-like, spherical, knobby or club-shaped swellings [12]. The root galls significantly alter the morphology, development, and physiology of the diseased plants [13]. Moreover, *P*. *brassicae* consumes carbohydrates from the viable cells in gall, and make gall into a sink for nutrient substances [14,15].

Tumourous stem mustard (*Brassica juncea* var. *tumida*) is an economical and nutritionally important vegetable crop widely grown in Fuling County, a district in Chongqing Municipality, China. Clubroot is one of the most harmful diseases of tumourous stem mustard causing substantial economic damage [16]. The diseased tumourous stem mustard shows small plants and swollen roots and finally dies due to root rot during the late plant growth stage in the field, although *P*. *brassicae* is an obligate parasite. Moreover, many pathogenic fungi such as *Fusarium* sp. are isolated from the clubroots, which indicates that the community of fungal endophyte may change during the infection of *P*. *brassicae*. However, the specific alteration in the community of fungal endophyte is still unclear.

In this context, the objectives of this study are, (1) to examine the diversity and composition of the fungal endophyte community in the roots of tumourous stem mustard (2) to demonstrate the changes in the community of fungal endophyte in the clubroots of tumourous stem mustard caused by *P*. *brassicae* compared to healthy roots.

## Materials and methods

### Sample collection

*P. brassicae*-infected tumourous stem mustard roots were obtained from three fields (February 2, 2017) in Fuling (29.21° N, 106.56° E) where clubroot disease was observed during the last 20 years. Roots were collected from the plants at the harvest-stage. The roots were classified as either R (healthy roots, no clubroot symptoms) or C (diseased roots, swollen clubroot symptoms). From one field, 30 plants including 15 R and 15 C samples were randomly selected and formed as one group. By this, three groups contained 90 plants from three fields and were referred to R1, C1, R2, C2, R3, and C3. The roots were washed with tap water to remove the soil particles. The healthy roots with a diameter of 0.5 cm from undiseased plants and clubroot galls with a diameter of 1 cm from diseased plants were cut off, and then sterilized by 70 % (v/v) ethanol for 40 s, followed by 4 % (w/v) sodium hypochlorite for 60 s, and finally washed three times using sterile distilled water. The surface sterilized healthy roots and galls peeled with a sterilized razor were divided into three portions. One portion was used to extract genomic DNA by cetyltrimethylammonium ammonium bromide (CTAB) DNA extraction method. The concentration and purity of DNA were monitored on 1% agarose gel. One portion was made into paraffin section, dyed with sarranine and observed under an optical microscope. Another portion was used to isolate endophytic fungi by the culture-dependent method.

### Isolation of endophytic fungi and its taxonomic identification

Three types of media, i.e. potato dextrose agar (BD Difco), rose bengal medium (BD Difco), and Czapek medium (BD Difco) were used for the isolation of endophytic fungi. The surface sterilized healthy roots, or clubroot samples were cut into tissue blocks of 5 mm x 5 mm, and were planted on a medium at 25 °C in the dark. When mycelia appeared, they were transferred into another medium for purification. A total of 156 fungal isolates were then identified based on the ITS sequence data. Mycelia of the fungal isolates were ground with liquid nitrogen in a sterile mortar, and Genomic DNA from all the fungal isolates was extracted using DNA extraction kit (TIANGEN Co. Ltd., Beijing, PR China). The internal transcribed spacer (ITS) region was amplified with universal primers (ITS1 and ITS4) [17]. PCR mixture contained 12 μL of 2× Taq PCR Mix (TIANGEN Co. Ltd., Beijing, PR China), 1 μL DNA template, 1 μL of each primer and 8 μL double distilled water. The PCR reaction conditions were as follows: initial pre-heating at 94 °C for 3 min, 30 cycles of 94 °C for 30 s, 58 °C for 30 s, 72 °C for 30 s, and a final extension at 72 °C for 10 min. PCR products were sequenced by 3730 sequencer (Majorbio Tech Co. Ltd., Beijing, PR China). The ITS sequence was aligned in the NCBI database (http://www.ncbi.nlm.nih.gov/) using the BLAST program. The highest hits (identity values higher than 97%) were regarded as the taxonomy of fungal isolates. The relative abundances were calculated according to the taxonomy of fungal isolates.

### High-throughput ITS Sequencing

ITS fragments were used as marker genes for high-throughput sequencing, and the primers of ITS fragments target most fungal groups, as well as other eukaryotes. The ITS fragments were amplified by PCR for barcoded pyrosequencing using the primers ITS1 and TS2 [18]. The employed PCR conditions were: 95 °C for 2 min (one cycle), 95 °C for 30 s, 55 °C for 30 s, and 72 °C for 30 s (25 cycles), and 72 °C for 5 min (one cycle). The sequencing was performed using an Illumina MiSeq sequencer (Asbios Technology Co., Ltd, China).

### Processing of Bioinformatics and Data Analysis

For diversity analyses, the bioinformatics analysis pipeline was conducted on the free online platform, Majorbio I–Sanger Cloud Platform (http://www.i-sanger.com). First, the raw sequences were processed using the QIIME package (v1.8) [19]. The low-quality sequences, such as primer and barcode sequence mismatch, and PCR-based or sequencing errors and chimeras, were removed. The taxonomy of each operational taxonomic unit (OTU) representative sequence was performed using Unite (Release 7.2) and ITS of *P. brassicae* under the threshold of 97% identity [20,21]. The Shannon index, Simpson index, and rarefaction curves were calculated to evaluate the α-diversity. Relative abundance of endophyte was calculated at phylum, genus and OTU levels. Analysis of variance (ANOVA) of the Shannon and Simpson indices was obtained by SPSS 16.0 and the relative abundance was used to assess the differences in the endophyte communities between the healthy roots and clubroots. For the evaluation of β-diversity, hierarchical cluster dendrograms (Bray-Curtis distance dissimilarities) were constructed according to the composition of OTU [22]. An unweighted UniFrac principal component analysis (PCA) and Principal coordinates analysis (PCoA) were performed using R 3.1.1 statistical software [23]. The differentially abundant genera between the healthy roots and clubroots were identified for finding the biomarker by Discriminant Analysis Effect Size (LEfSe) software [24]. Co-occurrence network analysis was conducted to reveal the relationship among the top 30 OTUs within the endophyte communities by Networkx software [25]. Pearson’s rank correlation coefficients between OTUs were calculated, and only the correlations with r > 0.5 were selected to construct the network.

## Results

### Symptoms of Clubroot and Detection of *P*. *brassicae*

Healthy roots were not swollen, and the lateral roots appeared normal, whereas clubroots were swollen and few lateral roots were observed (Fig. 1A). Abundant resting spores were found to be in the cells of clubroots (Fig. 1B), but no resting spores were observed in the healthy roots (Fig. 1C).

**Fig 1.**
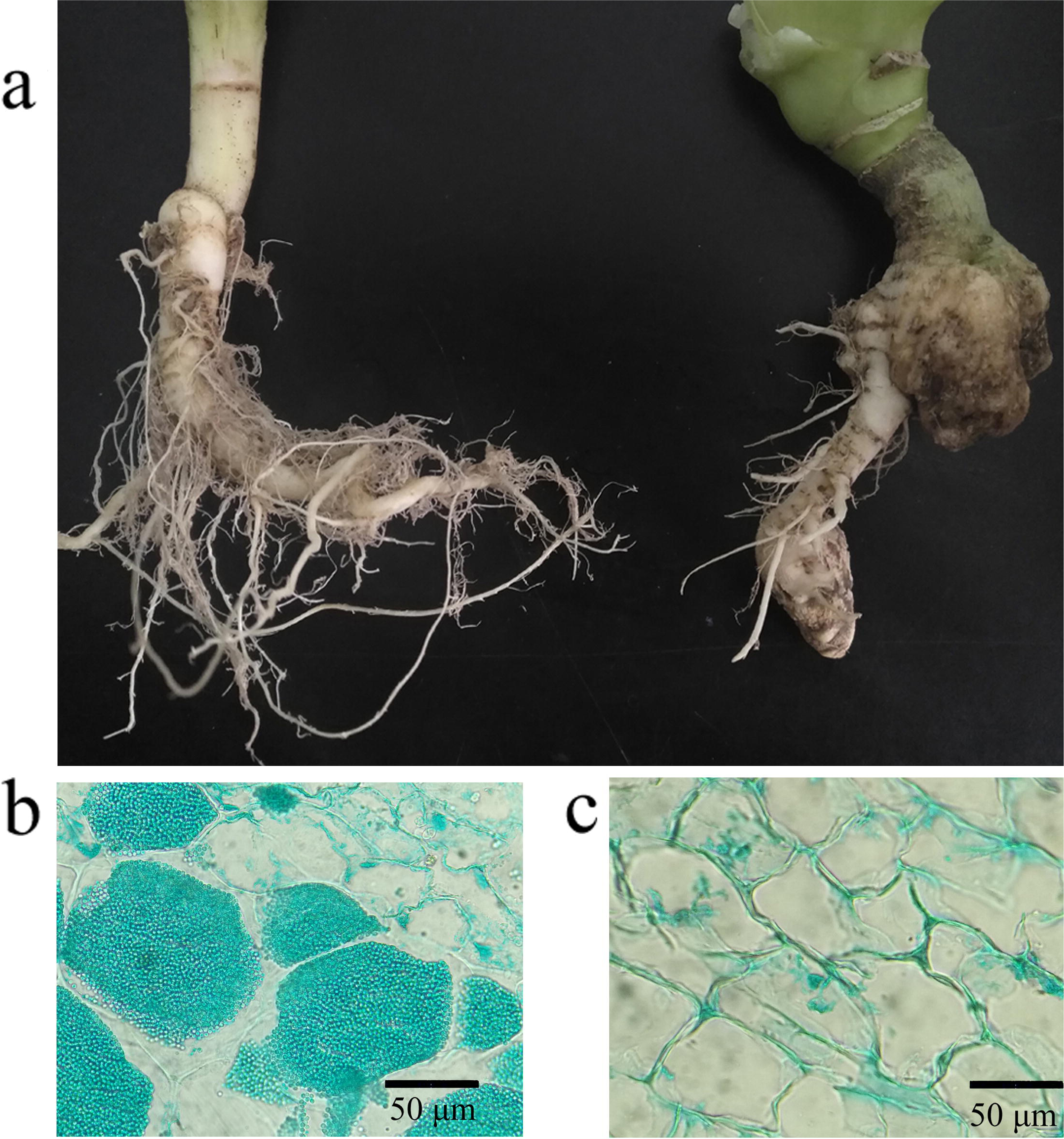
Healthy roots and clubroots of tumourous stem mustard caused by the infection of *P. brassicae* (A) roots of a healthy plant (left) and a diseased plant (right) (B) resting spores (pots) in the root cells of cluboorts coloured with fast green (C) the cell of healthy roots.

### Analysis of α-diversity

High-quality sequences of ITS were produced by the Miseq platform. The raw sequencing data for ITS sequencing were deposited at the Sequence Read Archive (SRA, https://www.ncbi.nlm.nih.gov/sra) under an accession number, SRP136514. The statistical formation of the sequences is shown in Table S1. According to the taxonomy of the sequence and abundance (Table S2), the composition of the endophyte communities was analyzed. Rarefaction curves analysis confirmed that the number of observed OTUs increased asymptotically with an increase in the reads (Fig. 2A). Both a higher Shannon index and Simpson index of the endophyte community were observed in the healthy roots than that in the clubroots (Fig. 2B), suggesting a higher diversity of the endophyte community in the healthy roots.

**Fig 2.**
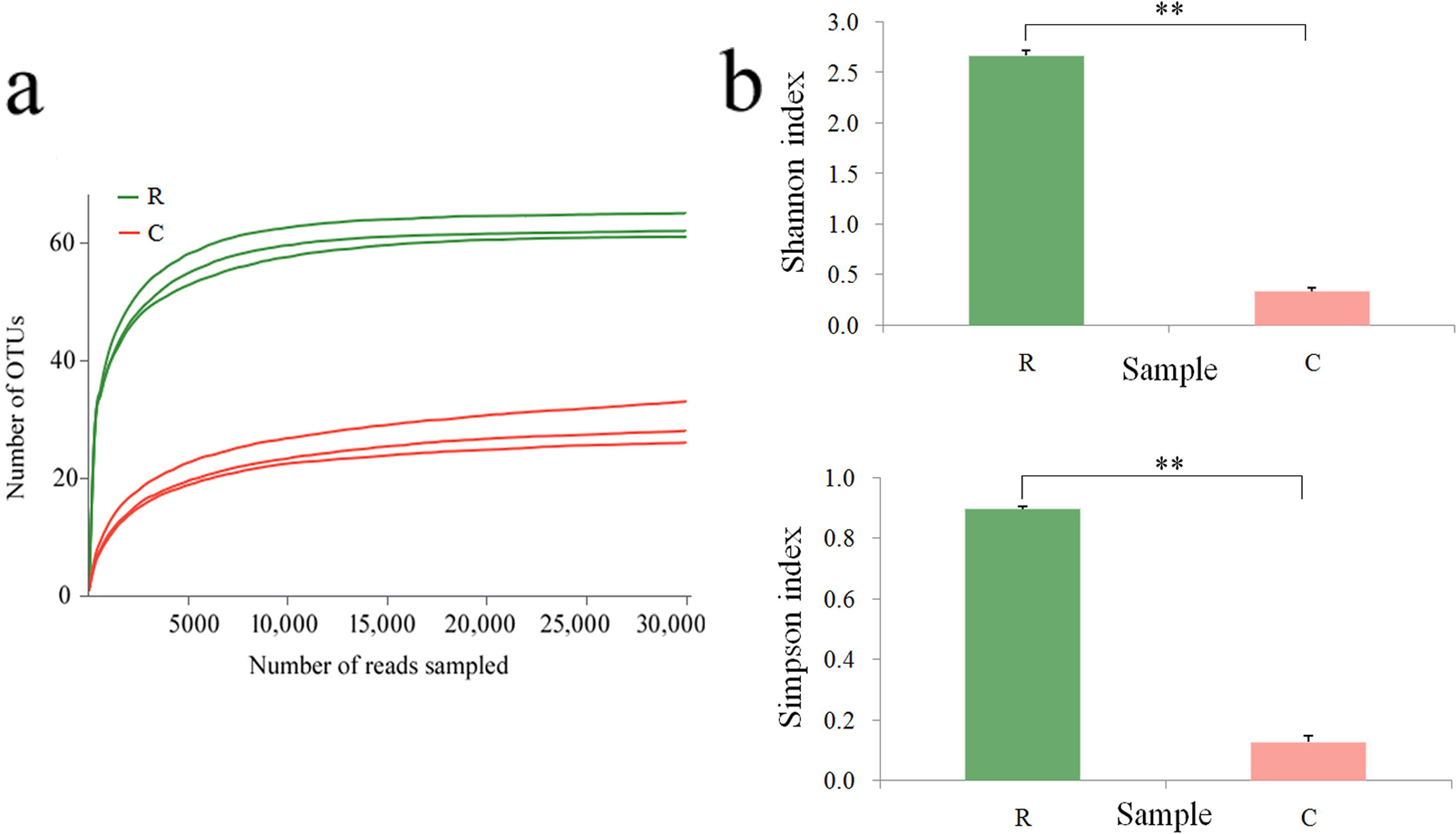
Rarefaction curves (A) and diversity indices (B) of the endophyte communities associated with the healthy roots and clubroots of tumourous stem mustard infected with *P. brassicae*. R, healthy roots. C, clubroots. ** differences at 0.01 level.

Ascomycota was the dominant taxon (a relative abundance ranging from 76.3 to 78.5%) of the endophyte community in the healthy roots followed by Zygomycota (RA, 15.5 to 19.9%), while Cercozoa (RA, 92.0 to 94.3%) was the most dominant taxon of the endophyte community in the clubroots followed by Ascomycota (RA, 5.6 to 7.6%) (Fig. 3A). The abundance of Basidiomycota and unclassified endophytic fungi was higher in the healthy roots compared to clubroots.

**Fig 3.**
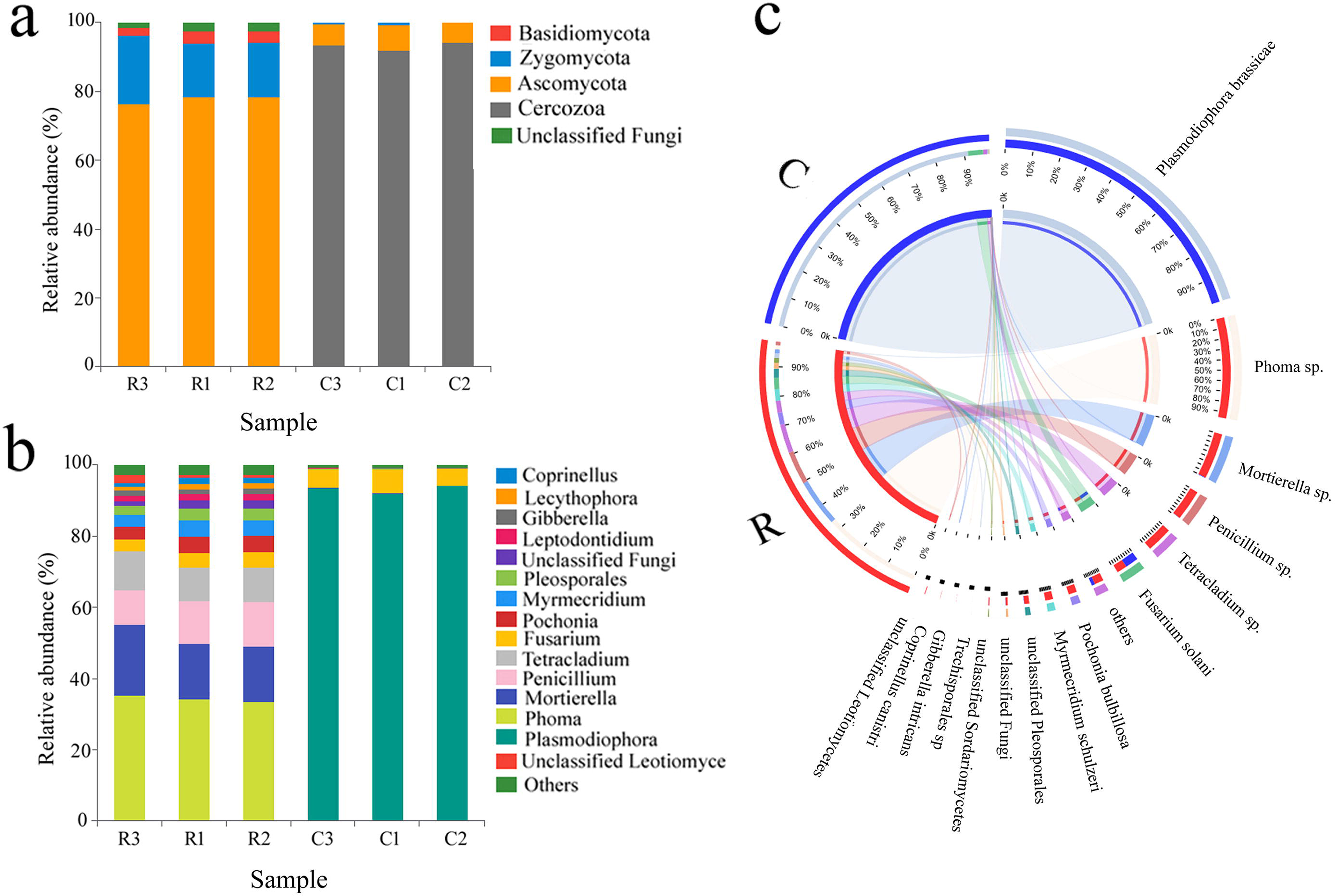
Relative abundance (%) of the endophyte at phylum (a), genus (B) and Circos figure at the genus level (C) in the healthy roots and clubroots of tumourous stem mustard infected with *P*. *brassicae*. R, healthy roots. C, clubroots.

At the genus level, a total of 40 and 27 genera were detected in the healthy roots and clubroots, respectively. In the healthy roots, *Phoma* was the dominant genus with an RA of 19.15%, followed by *Mortierella* (17.09%), *Penicillium* (11.49%), *Pseudallescheria* (10.75%), *Tetracladium* (9.98%), and *Fusarium* (7.88%) (Fig. 3B). In the clubroots, *Plasmodiophora* was the dominant taxon (93.28 %) followed by *Fusarium* (5.52 %) (Fig. 3B). Circos figure confirmed that *Plasmodiophora* was the most abundant and was found only in the clubroots, whereas *Phoma, Mortierella*, and *Penicillium* were predominant in the healthy roots (Fig. 3C).

A total of 66 and 40 OTUs were observed in the healthy roots and clubroots, respectively. Heatmap showed that OTU268 was dominant in the endophyte community associated with clubroots, whereas OTU106, OTU153, OTU192, OTU197, OTU114, OTU126, and OTU185 dominated the healthy roots (Fig, S1).

### Analysis of β-diversity

Hierarchical clustering analysis based on Bray-Curtis distance dissimilarities revealed that the endophyte communities in the healthy roots and clubroots clustered in the two branches (Fig. 4A). UniFrac-weighted PCA based on the composition of OTU showed variations between the healthy roots and clubroots with the first two axes indicating 81.08 and 11.71 % of the total variation (Fig. 4B). PCoA plot also clearly demonstrated similar results of PCA with the first two axes showing 99.34 and 0.62% of the total variation (Fig. 4C). The endophyte community in the healthy roots was clustered on the right side of the PCA and PCoA plot while the endophyte community in the clubroots was clustered on the left side, indicating a clear separation between the endophyte community in R and C samples.

**Fig 4.**
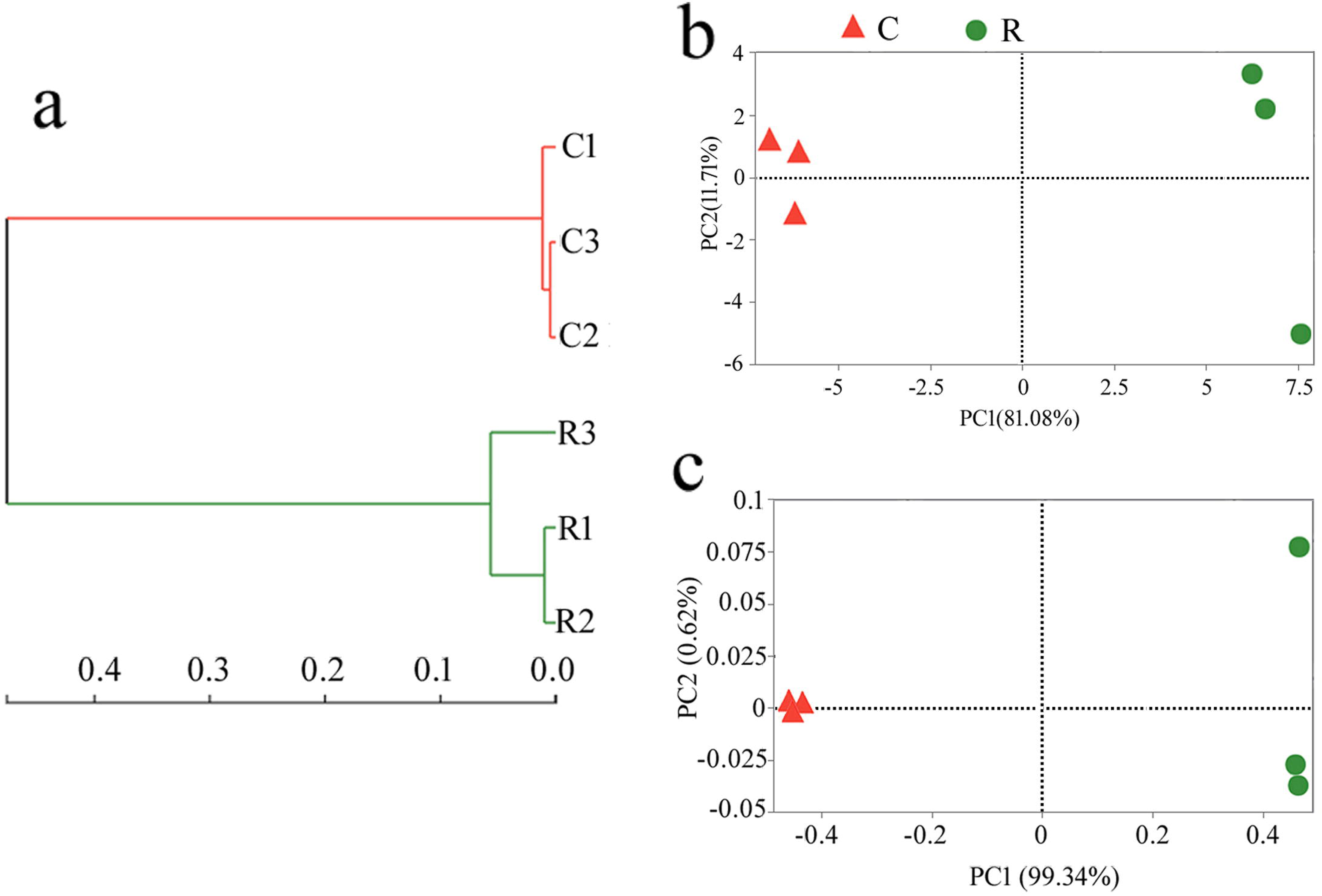
Hierarchical clustering analysis (a) UniFrac-weighted PCA (B) and PCoA (C) of the endophyte communities associated with the healthy roots and clubroots of tumourous stem mustard infected with *P*. *brassicae*. R, healthy roots. C, clubroots.

From the results of LEfSe, a significantly different taxa were found out between the two communities. At the genus level, *Phoma, Mortierella, Penicillium*, unclassified Pleosporales, etc. were enriched in the healthy root samples (Fig. 5A) and abundant *Plasmodiophora* in the clubroot samples possessed high values of linear discriminant analysis (LDA) (Fig. 5B). These taxa with different abundance can be regarded as potential biomarkers (LDA>3, P<0.05). Furthermore, 15 most abundant genera of the two endophyte communities were compared by Student’s t-test, where significant differences were observed (Fig. S2). For instance, *Plasmodiophora* was significantly more abundant in the clubroots while all other genera except *Fusarium* were significantly more abundant in the healthy roots.

**Fig 5.**
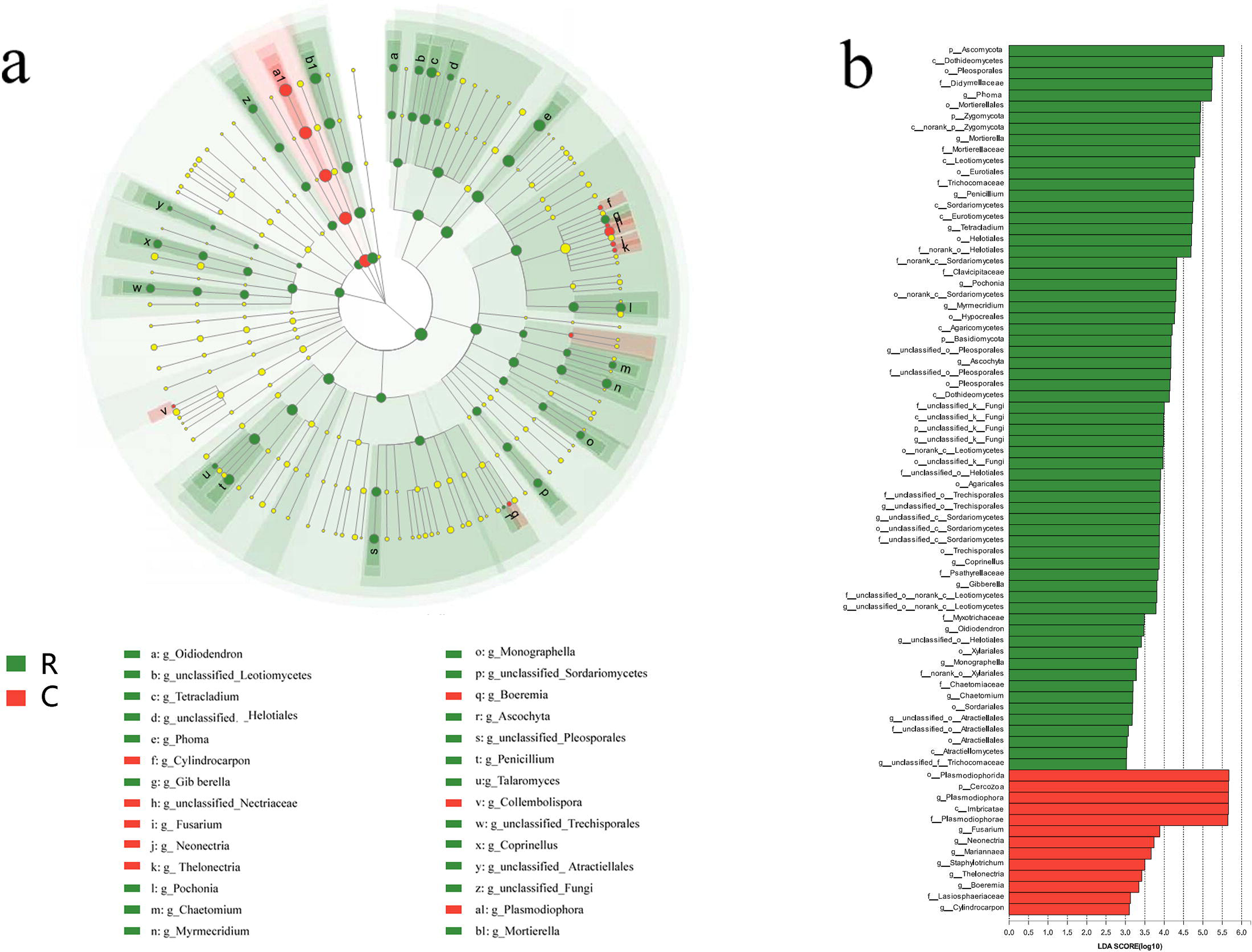
LefSe analysis (A) The cladogram diagram shows the taxas with marked differences in the two endophyte communities. Red and green indicate different groups, with the classification of taxas at the level of phylum, class, order, family, and genus shown from inside to the outside. The red and green nodes in the phylogenetic tree represent taxas that play an important role in the two endophyte communities, respectively. Yellow nodes represent taxas with no significant difference. (B) Species with the significant difference that have an LDA score higher than the estimated value; the default score is 3.0. The length of the histogram represents LDA score; i.e., the degree of influence of taxas with a significant difference between different groups. R, healthy roots. C, clubroots.

### Network Analysis

The endophyte community in the healthy roots had two centers with a complex network structure (28 nodes and 336 edges) (Fig. 6A) while the endophyte community in the clubroots had three centers with a less complex network structure (18 nodes and 161edges) (Fig. 6B). Most of the nodes belong to Ascomycota in both the healthy roots and clubroots networks and had the highest number of correlations with other taxa. The node of the Cercozoa (i.e., *Plasmodiophora*) only appeared in the clubroot network and had the highest percentage. When the relationship between *Plasmodiophora* and other fungi was analyzed, it was found that 10 OTUs had a significant correlation with *Plasmodiophora* (3 OTUs had a positive correlation, and 7 OTUs had negative correlation) (Table S3).

### Culturable endophytic fungi

The same samples for Miseq sequencing were also used to isolate endophytic fungi by the culture-dependent method. At the genus level, *Mortierella, Sarocladium, Phoma, Penicillium, Plectosphaerella*, and *Fusarium* were obtained (Fig. 7A). *Fusarium* (a relative abundance of 81.4%) was the predominant group in the clubroots, among which the following were present: *F. graminearum* (RA, 5.5 %), *F. oxysporum* (RA, 28.8 %), *F. solani* (RA, 24.0 %), *F. asiaticum* (RA, 8.2 %), and *Fusarium* sp. (RA 14.4 %). In the healthy roots, *Mortierella* was the main group with an RA of 36.2% followed by *Phoma, Penicillium, Fusarium*, etc. (Fig. 7B).

**Fig 6.**
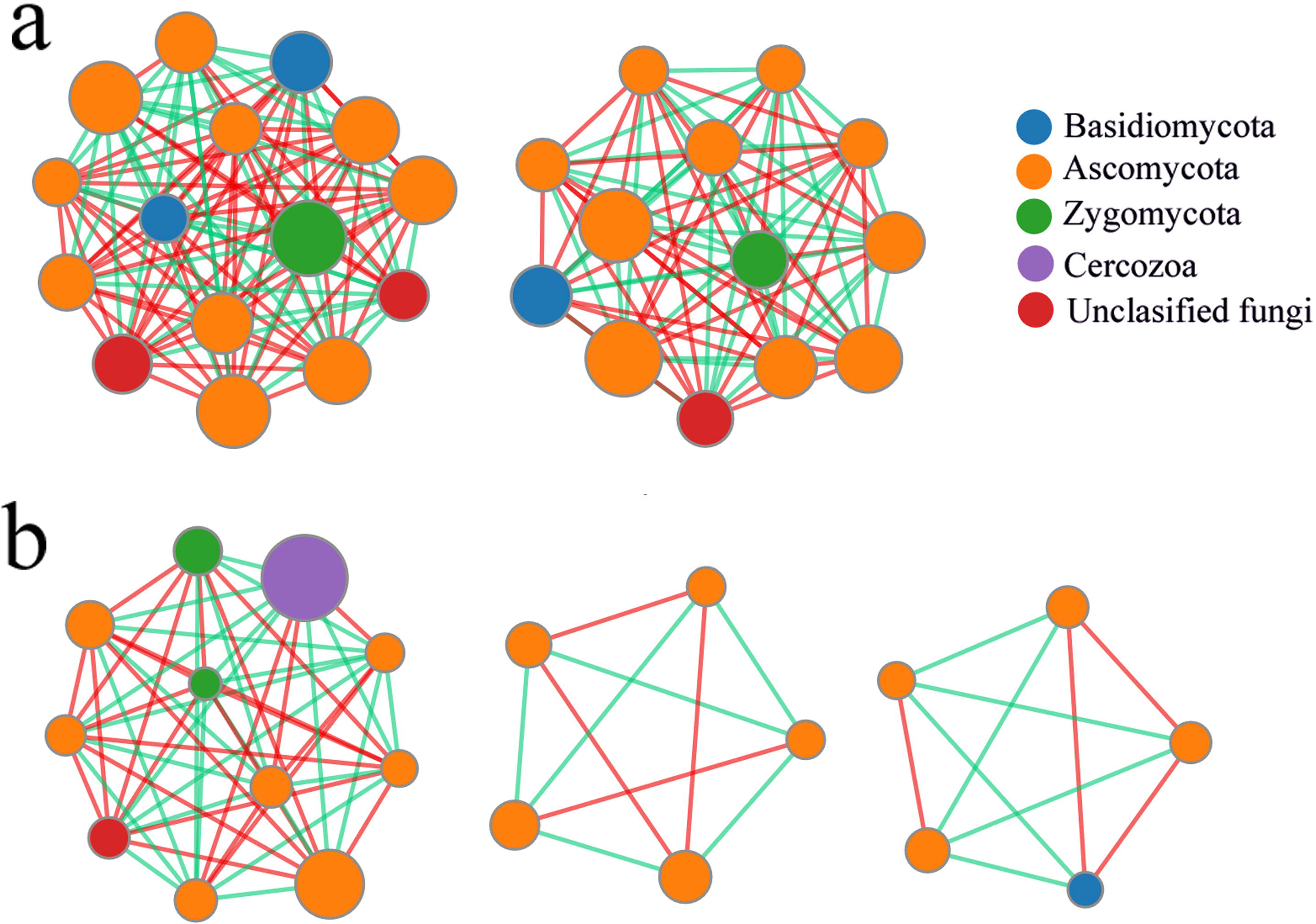
Co-occurrence network analysis of the two endophyte communities within the healthy roots and clubroots of tumourous stem mustard infected with *P. brassicae* (A) Healthy roots (B) Clubroots. Each node represents taxa affiliated at the OTU level, and the size of the nodes represent an average abundance of OTU. The lines represent the connections between each OTU. A red line indicates a positive correlation. whereas a green line indicates a negative correlation.

**Fig 7.**
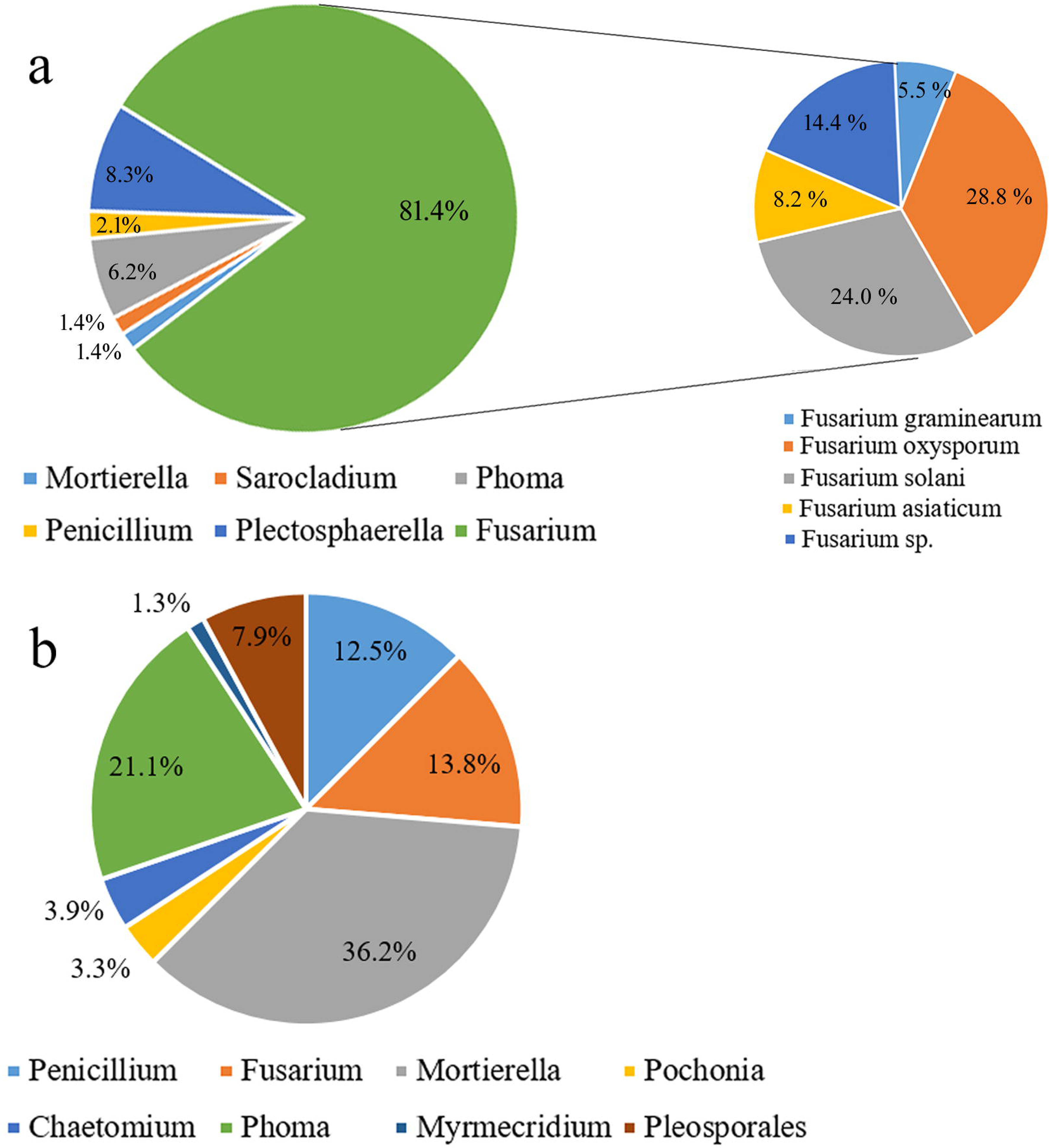
The endophytic fungi isolated from the clubroots (A) and healthy roots (B) by the culture-dependent method.

## Discussion

In this study, a high diversity of fungal endophyte in the healthy roots of tumourous stem mustard has been found out using high-throughput sequencing. Our results agree with the early reports that the plants can harbor a diversity of endophytic fungi in their healthy tissues, especially in the root system which is considered to be the most suitable habitat for endophytic fungi [26,27]. Ascomycota is reported to be the most common group of endophytic fungi [28], suggesting that they are suitable for the ecological niche of plant tissue. Zhao also reported Ascomycota as the dominant group of endophytic fungi in the roots of oilseed rape (*Brassica napus*) [29]. Also in this study, Ascomycota has been found to be the dominant taxon in the healthy roots of tumourous stem mustard. In the phylum Ascomycota, *Phoma, Mortierella, Penicillium*, and *Fusarium* were the common fungal genera in the healthy roots of tumourous stem mustard. Arie et al. reported that *Phoma glomerata* produces epoxydon and inhibits clubroot disease, demonstrating that *Phoma* in the healthy roots of tumourous stem mustard may perform a similar function [30,31]. *Mortierella* was also detected in the roots of oilseed rape using high-throughput sequencing [29]. Melo et al. confirmed that the endophytic *M. alpina* in the moss Schistidium antarctici produces antibiotics, antioxidant substances, and polyunsaturated fatty acids, which improve the environmental suitability of the plant host [32]. Wani et al. also found that endophytic *M. alpina* promotes the biosynthesis of Crocus apocarotenoid and enhances environmental stress tolerance [33]. These results reflect that the endophytic *Mortierella* is a benefit to the plant, which indicates that *Mortierella* in tumourous stem mustard is profitable to the host. Marinho et al. and Lin et al. both reported that endophytic *Penicillium* produces active polyketides [34,35], reflecting that *Penicillium* in tumourous stem mustard may produce similar substances.

Worldwide *Fusarium* is considered to be one of the most ubiquitous groups of fungi which can survive in a wide range of plants, environments, and climates [36]. In this study, some OTUs identified as *Fusarium* had a relatively high abundance in both the healthy roots and clubroots, reflecting the dominant presence of *Fusarium* in tumorous stem mustard. Previous studies show that most of the *Fusarium* species are nonpathogenic while only a small number are pathogenic [37]. Some OTU identified as *Fusarium* in the healthy roots perhaps were nonpathogenic because of the absence of wilt symptoms. OTU244 and OTU287 assigned as *F. solani*, a common soil-borne pathogen, were only and mainly found in the clubroots and these populations may be pathogenic. Moreover, *F. solani* is the most fungal isolate obtained by the culture-dependent method (Fig. 7A). Also, other pathogenic *Fusarium* such as *F. graminearum, F. oxysporum*, and *F. asiaticum* also isolated from the clubroots rather than the healthy roots, suggesting that *Plasmodiophora* infection many facilitate *Fusarium* to enter into the diseased roots. The presence of a plenty of pathogenic *Fusarium* in the clubroots maybe the reason for the death of tumourous stem mustard during the late plant growth stage in the field. In clubroots, *Plasmodiophora* was the extremely abundant genus, although not belonging to fungi, amplified by PCR with ITS primers for fungi and other eukaryotes. Zhao et al. also found that *Plasmodiophora* mainly enriched in the clubroots of *Brassica napus* by high throughput sequencing using the same primers [29].

In general, the healthy plant tissues harbor a more diverse community of endophytic fungi than diseased plant tissues [29,38]. Also, the endophyte community in the healthy roots of tumourous stem mustard had significantly higher Shannon and Simpson indices compared with the clubroots. Also, a higher number of genera and OTUs were obtained from the healthy roots than from the clubroots. The endophyte communities in the healthy roots and clubroots also differed in the α- and β-diversity. The marked difference in the composition of the endophyte community at the genus level between the healthy roots and clubroots as revealed by LEfSe and Student’s t-test showed that the dominant genera as biomarkers caused differences in the two communities. These substantial discrimination in the dominant genera was derived from the infection of *P. brassicae*. Pathogens may compete with the endophytic fungi for space and nutrients in the same niche within the plant tissue [39]. We presume that a decrease in the diversity of the fungal endophyte community in the clubroots of tumourous stem mustard has two causes. First, the physiological changes and the formation of clubroots induced by *P. brassicae* may affect the availability of nutrients for endophytes. Second, in clubroots, the resting spores of *P. brassicae* are produced in the galls of diseased plants and may fill the gall cells. This may restrict the available space for the endophytes. The phenomenon of a decrease in the diversity of endophytic fungi has been observed in many diseased plant species [29,40,41].

In this study, some plant pathogenic fungi were detected from the clubroots, such as *F. solani*, but not from the healthy roots of tumourous stem mustard. Similarly, Zhao et al. isolated many soil-borne pathogenic fungi, including *Fusarium, Gibberella, Alternaria, Sclerotinia, Leptosphaeria*, and *Cylindrocarpon*, from the oilseed rape roots infected with *P. brassicae* [29]. This multi-species aggregation involving *P. brassicae* and other pathogenic fungi is similar to disease complexes containing plant-parasitic nematodes and soil-borne pathogens [42]. The cuticle of clubroots is cracked, thereby providing entry points for pathogens, even for opportunistic pathogens. Additionally, the cracked cuticle aggravates the leaking of nutrients, such as amino acids and carbohydrates, into the rhizosphere soil which may attract soil-borne pathogens and increase the risk of infection by these pathogens [43]. Root-knot nematodes may form disease complexes with *Fusarium* species such as *F. oxysporum* and *F. solani* [44]. In this study, the abundant presence of *F. solani* in the clubroots indicates that this pathogen and *P. brassicae* may form a disease complex, which may make it more difficult to control the clubroot disease. Further studies are necessary to clarify the potential existence of this disease complex. Besides, fungi such as *C. cupreum, Phoma* sp. *Pochonia* sp. and *Myrmecridium schulzeri* were harvested and were reported as biocontrol fungi [45-47], reflecting that these fungi may help the host to resist the infection of *P. brassicae*.

Endophytic fungi in the plants must deal with many interactions and build a balanced network [48]. In this study, the endophyte community network in the healthy roots of tumourous stem mustard was observed to be more complicated than in the clubroots, indicating a balanced network of interactions among the fungal endophyte community in the healthy roots. In the clubroots, *P. brassicae* dominated the endophyte community which was out of balance showing a weak network of interactions. In general, endophytes are more diverse and abundant than the pathogens in a healthy plant microbiome, and *vice versa* in a diseased plant microbiome [49].

## Conclusions

In conclusion, the obtained data from this study showed that the fungal endophyte community in the clubroots are markedly different from the healthy roots in terms of alpha and beta diversity, suggesting that the infection of *P. brassicae* changes the fungal endophyte community in tumourous stem mustard roots. Future work should involve identifying the pathogenicity of *F. oxysporum* and *F. solani* on tumourous stem mustard. Moreover, some endophytic fungi with biocontrol activity against *P. brassicae* could also be considered in the evaluation.

## Supporting information

**Table S1 Sequence information of each sample**. R, healthy roots. C, clubroots.

**Table S2 Taxonomy and distribution of the OTUs**. Taxonomy at Phylum, Class, Order, Family, Genus, Species, and OTU level. R, Healthy roots. C, Clubroots. The number in the table cell is the number of sequences of each OTU.

**Table S3 Pearson’s correlation relationships between *Plasmodiophora* and OTUs showed in the network**.

**Fig S1 Heatmap of the 50 most abundant OTU from R and C samples**. R, healthy roots; C, clubroots.

**Fig S2 Student’s t-test bar plot of the endophyte communities at the genus level in the healthy roots and clubroots of tumourous stem mustard infected with *P*. *brassicae***. p<0.05*, p <0.01**, p<0.001***. R, healthy roots. C, clubroots.

