## Supplement Table 1 for "Infection of *Plasmodiophora brassicae* changes the fungal endophyte community of tumourous stem mustard roots as revealed by high-throughput sequencing and culture-dependent methods"

**S1Table Sequence information of each sample**

| Sequence information | Sequence number | Mean length | Minimum length | Maximum length |
| --- | --- | --- | --- | --- |
| R1 | 38688 | 277.035 | 221 | 390 |
| R2 | 60474 | 276.712 | 216 | 390 |
| R3 | 30448 | 274.834 | 200 | 356 |
| C1 | 33749 | 201.849 | 180 | 285 |
| C2 | 42868 | 200.842 | 181 | 289 |
| C3 | 31825 | 201.078 | 180 | 310 |

R: healthy roots. C: clubroots
