## Supplement Table 3 for "Infection of *Plasmodiophora brassicae* changes the fungal endophyte community of tumourous stem mustard roots as revealed by high-throughput sequencing and culture-dependent methods"

**S3 Table** Pearson’s correlation [relation](app:ds:relation)ships between *Plasmodiophora* and OTUs showed in network.

| Taxonomy | OTU | Correlation index |
| --- | --- | --- |
| unclassified Pleosporales | OTU123 | -0.97 |
| unclassified Mortierella | OTU153 | -0.90 |
| Wallemia mellicola | OTU107 | -0.88 |
| unclassified Fusarium | OTU185 | -0.78 |
| unclassified Sordariomycetes | OTU144 | -0.40 |
| s__unclassified_k__Fungi | OTU235 | -0.23 |
| Pyrenochaetopsis leptospora | OTU246 | 0.47 |
| Gibberella intricans | OTU52 | 0.50 |
| Monographella cucumerina | OTU176 | 0.61 |
| Fusariumsolani | OTU287 | 0.73 |
| Gibberella zeae | OTU319 | 0.76 |
| unclassified Gibberella | OTU282 | 0.78 |
| Nectria ramulariae | OTU310 | 0.88 |
| Phoma macrostoma | OTU294 | 0.97 |
| Metarhizium marquandii | OTU321 | 0.99 |

Metagenomic biomarker discovery and explanation
