## Supplementary figures and images for "Infection of *Plasmodiophora brassicae* changes the fungal endophyte community of tumourous stem mustard roots as revealed by high-throughput sequencing and culture-dependent methods"

### Supplement Figure 1

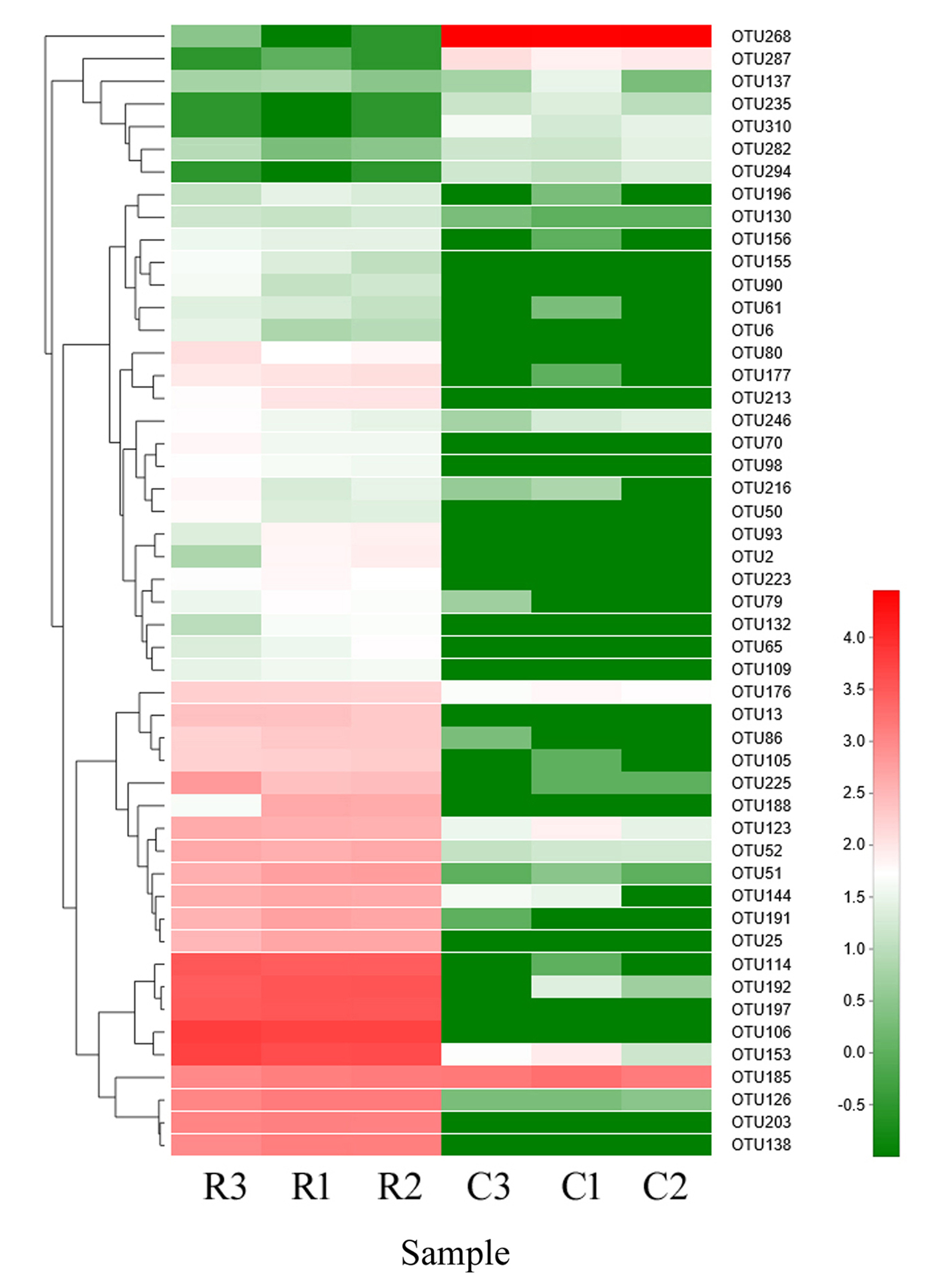

### Supplement Figure 2

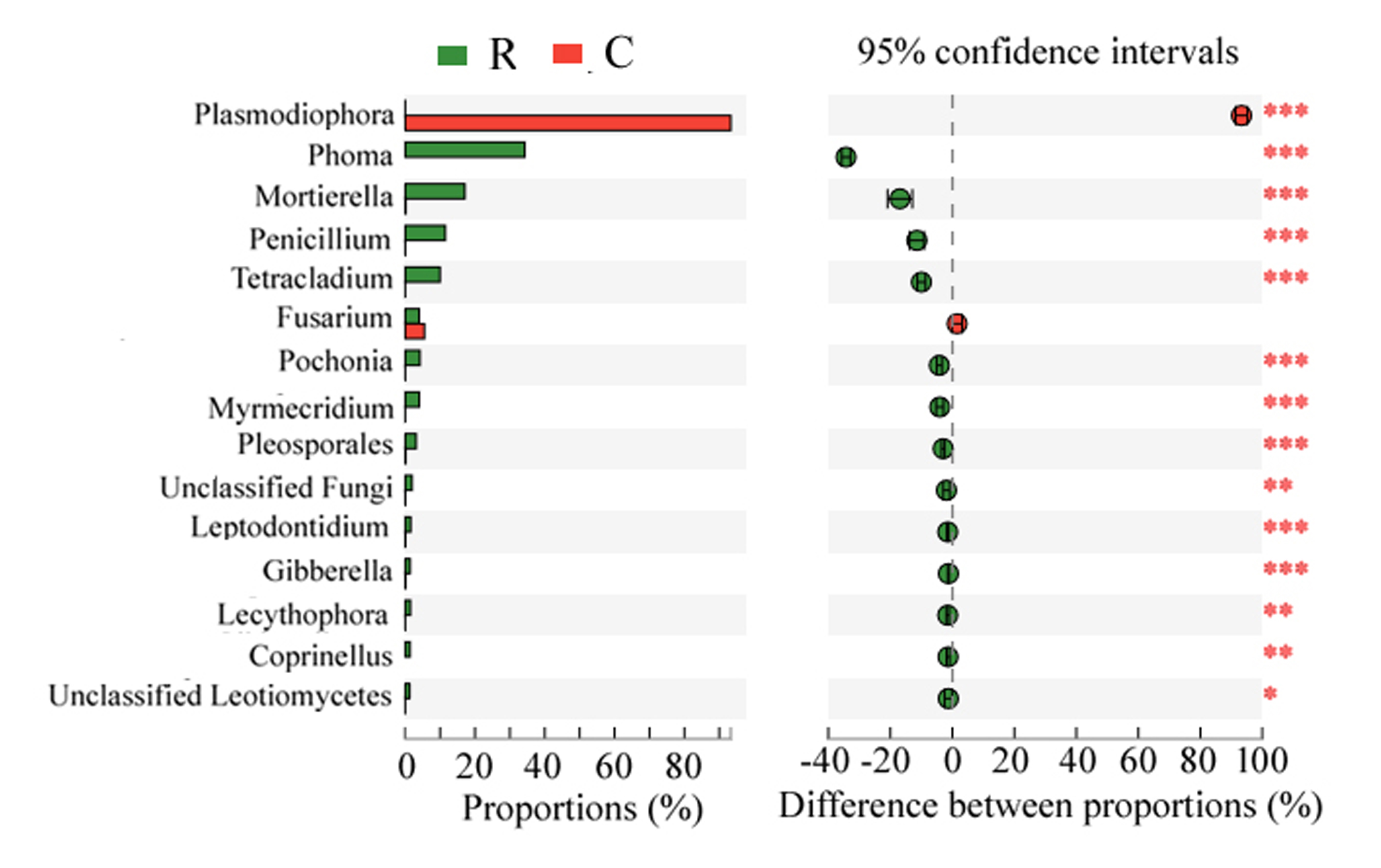
